# Ovarian cancers with low CIP2A tumor expression constitute an APR-246 sensitive disease subtype

**DOI:** 10.1101/2021.03.31.437804

**Authors:** Anna N. Cvrljevic, Umar Butt, Kaisa Huhtinen, Tove J. Grönroos, Camilla Böckelman, Heini Lassus, Katja Kaipio, Tiina Arsiola, Teemu D. Laajala, Denise C. Connolly, Ari Ristimäki, Olli Carpen, Jeroen Pouwels, Jukka Westermarck

**Author notes:** The authors have declared that no conflict of interest exists.

## Abstract

Identification of ovarian cancer (OvCa) patient subpopulations with increased sensitivity to targeted therapies could offer significant clinical benefit. We report that 22% of the high grade OvCa tumors at diagnosis express CIP2A oncoprotein at low levels. CIP2A^low^ OvCa tumors have significantly lower likelihood of disease relapse after standard chemotherapy, but yet a portion of relapsed tumors retain their CIP2A^low^ phenotype. We further discover that reactive oxygen species (ROS) inducing compound APR-246 (PRIMA-1Met/Eprenetapopt), currently in clinical development, preferentially kill CIP2A^low^ OvCa cells across multiple chemotherapy resistant cell lines. Consistent with CIP2A^low^ OvCa subtype in humans, CIP2A is dispensable for development of MISIIR-TAg-driven mouse OvCa tumors. Nevertheless, CIP2A deficient OvCa tumor cells from MISIIR-TAg mice displayed APR-246 hypersensitivity both *in vitro* and *in vivo*. Mechanistically, the lack of CIP2A expression hypersensitizes the OvCa cells to APR-246 by inhibition of NF-kB activity. Accordingly, combination of APR-246 and Nf-kB inhibitor compounds strongly synergized in killing of CIP2A positive OvCa cells. Collectively, we discover low CIP2A expression as a vulnerability for APR-246 in OvCa. The results warrant consideration of clinical testing of APR-246 for CIP2A^low^ OvCa tumor subtype patients, and reveal CIP2A as a candidate APR-246 combination therapy target.

## Introduction

Ovarian cancer is the fifth most common cause of cancer-related death among females in the United States. In the United States alone, every year more than 22, 000 women receive OvCa diagnosis, and around 14, 000 women die from this disease. Although most patients with primary OvCa respond well to standard adjuvant chemotherapy, the 5-year disease-specific overall survival in OvCa has been historically less than 50%, and during progression the disease becomes resistant to most current therapies (1). However, as evidenced by a significant clinical benefit of poly ADP ribose polymerase (PARP) inhibitors for platinum-sensitive OvCa patients, identification of new therapies for patient subpopulations with enhanced therapeutic response, might significantly change the disease outcome of those OvCa patients (2).

Tumor suppressive protein phosphatase 2A (PP2A) complexes control activities of number of oncogenic proteins and cancer driver pathways (3). In many cancer types, the tumor suppressor activity of PP2A is suppressed by its endogenous inhibitor protein CIP2A (4, 5). CIP2A has a restricted expression profile in most human and mouse normal tissues (5, 6), but it is overexpressed with high frequency in most human malignancies (4, 7). High CIP2A expression has been observed in 68-83% of high-grade serous OvCa tumors, and this associates with high proliferation index, aneuploidy, advanced tumor grade, *TP53* mutation, and EGFR expression (8, 9). On the other hand, the remaining 17-32% of OvCa patients with CIP2A^low^ expressing tumors have significantly longer overall ovarian cancer-specific survival both in unselected patient population, as well as among patients treated with standard platinum-based chemotherapy (8). CIP2A was recently also shown in cell culture to protect OvCa cells from Cisplatin-induced apoptosis (10), and to associate with stemness features in patient-derived high grade serous cancer (HGSC) cells (11). Further, in two cancer drug response screens, CIP2A depletion was shown to increase therapeutic response of HeLa and KRAS-mutant lung cancer cells to various types of cancer therapies (12, 13). Together, these results indicate that CIP2A^low^ OvCa tumors, consisting of approximately 1/5 of all OvCa patients, may constitute a less aggressive, and more therapy sensitive OvCa subtype. The aim of this study was to identify clinically applicable compounds that would preferentially kill CIP2A^low^ OvCa cells. Discovery of such compounds could potentially provide basis for predictive patient stratification strategy for OvCa patients with CIP2A^low^ tumor subtype (2).

## Results and discussion

### Screening for therapeutics that preferentially kill CIP2A^low^ OvCa cells

In a previously described retrospective cohort of 562 serous OvCa patients treated with standard chemotherapy (8), and for which both CIP2A status by immunohistochemistry (IHC), and relapse status was known, 266 patients achieved complete response (CR) after treatment with surgery and 6-8 rounds of paclitaxel-carboplatin combination. Among this group, 21,4% of tumors had negative CIP2A protein expression (Table S1). Notably, patients with CIP2A negative OvCa tumors at diagnosis significantly more often achieved complete response (CR) than patients with CIP2A positive tumors (57% vs. 45%, chi squared test p-value 0,044). OvCa tumor CIP2A negativity also very significantly predicted lower likelihood for disease relapse after chemotherapy (Table S1).

These results indicate that a portion of OvCa tumors develop in a CIP2A-independent manner. Results also support the earlier findings that CIP2A^low^ tumors could constitute a more therapy sensitive subtype (10, 12, 13). To identify potential novel therapies for the CIP2A^low^ OvCa subtype, we conducted a drug screen comparing cell viability effects of clinically used, or experimental drugs, between CIP2A^high^ (control shRNA) and CIP2A^low^ (CIP2A shRNA) HEY cells. Inhibition of CIP2A expression was confirmed by Western blotting (Fig. S1A). CIP2A^high^ cells showed multi-drug resistance against chemotherapies commonly used used for OvCa (Cisplatin, Doxorubicin, Olaparib, Paclitaxel, Topotecan)(Fig. 1A). However, CIP2A^low^ HEY cells were at least to certain extent more sensitive to the majority of tested drugs at chosen concentrations (Fig. 1A). The most apparent sensitization effect was observed with APR-246 (PRIMA-1Met/Eprenetapopt) (14–18). Whereas, CIP2A^high^ cells were practically insensitive to APR-246, CIP2A^low^ HEY cells showed > 50% reduction in cell viability (Fig. 1A). APR-246 (Eprenetapopt) has been studied in clinical trial in OvCa (19), and it showed promising clinical activity in a recent phase II trial in acute myeloid leukemia (AML) (18).

**Figure 1.**
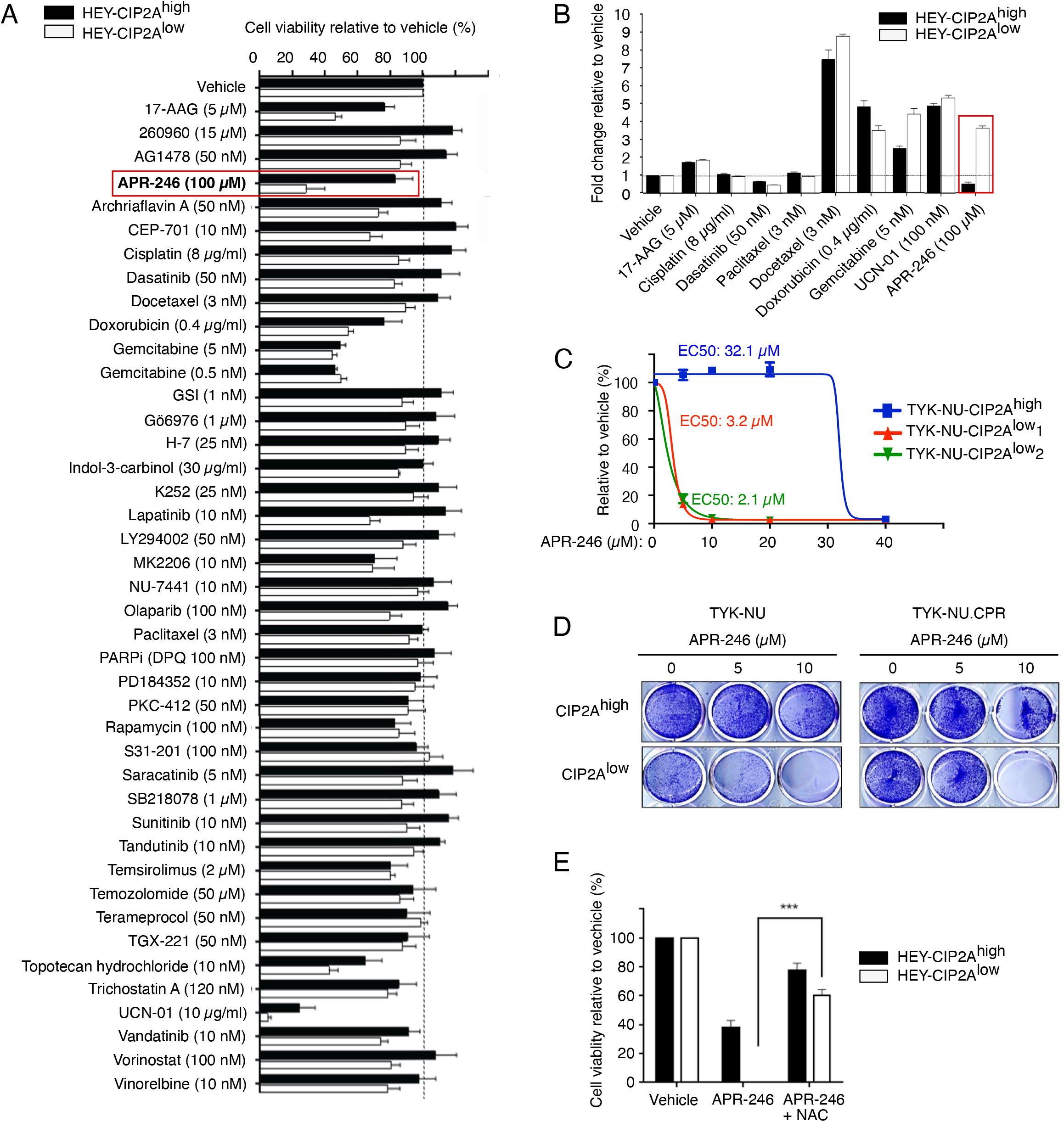
Identification of APR-246 hypersensitivity in CIP2A^low^ ovarian cancer cells. **A)** Relative cell viability of HEY cells stably transduced either by control shRNA (HEY-CIP2A^high^) or by CIP2A targeted shRNA (HEY-CIP2A^low^) treated with indicated cancer therapeutics for 24 hours. Shown is mean + S.D. from parallel samples from representative screen. **B)** Relative caspase 3/7 activity in HEY-CIP2A^high^ or HEY-CIP2A^low^ cells treated with indicated cancer therapeutics for 48 hours. Shown is mean + S.D. of parallel samples from representative screen. **C)** Relative cell viability of TYK-NU cells stably transduced either by control shRNA (TYK-NU-CIP2A^high^) or by two CIP2A targeted shRNAs (TYK-NU-CIP2A^low1,2^) treated with increasing concentrations of APR-246 for 48 hours. Shown is mean + S.D. of parallel samples from representative screen. EC50: half maximal effective concentration. **D)** Colony growth assay of TYK-NU-CIP2A^high^ and TYK-NU-CIP2A^low^1,2 cells and their cisplatin resistant derivatives (TYK-NU-CPR) treated with indicated doses of APR-246. **E)** Relative cell viability of HEY-CIP2A^high^ or HEY-CIP2A^low^ cells treated with either APR-246 (20 μM) alone or in combination with N-acetyl cysteine (NAC)(5 mM) for 48 hours. Shown is mean + S.D. of parallel samples from representative experiment. p<0.001.

To validate these results, and to understand the mode of cell killing by APR-246 in CIP2A^low^ HEY cells, we screened nine of the drugs by using caspase3/7 apoptosis assay. HEY cells were resistant to 17-AAG, Cisplatin, Paclitaxel, and Dasatinib, regardless of their CIP2A status (Fig. 1B). On the other hand, Docetaxel, Doxorubicin, Gemcitabine, and UCN-01 induced caspase3/7 activity in CIP2A^high^ cells. Notably, APR-246 was the only drug that did not induce apoptosis in CIP2A^high^ cells, but showed clearly higher apoptotic response in CIP2A^low^ cells (Fig. 1B). Apoptosis induction in APR-246 treated CIP2A^low^ cells was confirmed by COMET assay by using two independent shRNA sequences (Fig. S1A,B).

To confirm that vulnerability of CIP2A^low^ cells to APR-246 was not restricted to HEY cells, we tested the impact of CIP2A for APR-246 response in HGSC cell line TYK-NU. In a cell viability assay, CIP2A^low^ TYK-NU cells showed dramatically decreased EC50 values for APR-246 as compared to control shRNA expressing cells, and there was no difference between CIP2A^low^ cells expressing two independent CIP2A shRNA sequences (Fig. 1C). Hypersensitivity of CIP2A^low^ cells to APR-246 was also confirmed by colony growth assays in TYK-NU, and its cisplatin-resistant derivative TYK-NU.CPR cell line (Fig. 1D). To confirm that the effects were not related to clonal selection of shRNA transduced cells, and to expand the results to yet other OvCa cell lines, we transiently inhibited CIP2A expression by siRNA transfection in HEY, CAOV-3, NIH:OVCAR3, SKOV-3 and OVCAR-8 cells. In all cell lines CIP2A silencing resulted in increased sensitivity to APR-246 in a cell viability assay (Fig. S1C).

Although APR-246 was originally identified as a compound that reactivates mutant TP53 (15, 16, 20), the tested OvCa cell lines displaying hypersensitivity to APR-246 upon CIP2A inhibition, exhibit varying *TP53* mutation statuses. Whereas HEY is *TP53* wild-type, and SKOV-3 has both *TP53* alleles deleted, the rest of the cells lines harbor distinct *TP53* mutations: TYK-NU (R175H); NIH:OVCAR3 (R248Q); CAOV-3 (Q136*); and OVCAR8 (Y126_K132del; c.376-396del) (21) (https://p53.iarc.fr;https://web.expasy.org/cellosaurus/). Therefore, it is unlikely that the cell killing effects by APR-246 in the tested OvCa cells would be mediated solely by its mutant TP53 reactivating activity. On the other hand, several recent studies (using some of the same OvCa cells as here) have shown that APR-246 kills cancer cells independently of TP53, but via induction of reactive oxygen species (ROS) (16, 21, 22). Moreover, a recent study showed that MQ, the active product of APR-246 in cells, conjugates with GSH to disrupt the cellular antioxidant balance (17). In a similar vein, we observed APR-246-elicited induction of ROS production in HEY cells, and this was completely quenched by pre-treatment of cells with anti-oxidant N-acetyl cysteine (NAC) (Fig. S1D). Strongly supporting ROS induction as a causative mechanism for APR-246-elicited cell killing of CIP2A^low^ cells, NAC pre-treatment prevented the effects of APR-246 in cell viability (Fig. 1E). Of a note, the high micromolar concentrations of APR-246 required for OvCa cell killing is consistent with published studies (17, 21), and due to intracellular metabolism of the drug to the active product methylene quinuclidinone (MQ)(17). Further, experiments shown in figure 1A and 1B were performed with drug patch that apparently had lower bioactivity, and hence up to 100 uM concentrations had to be used, whereas the rest of the experiments were performed with APR-246 provided generously by APREA Therapeutics developing APR-246 (Eprenetapopt^®^) towards clinical cancer therapy.

Collectively, these results identify low CIP2A expression as a vulnerability to ARP-246 across multiple chemotherapy resistant OvCa cell lines.

### Low CIP2A expression confers OvCa cell APR-246 hypersensitivity *in vivo*

*In vivo* relevance of CIP2A on OvCa cell APR-246 sensitivity was assessed by subcutaneous xenograft assay with stable shRNA transduced HEY cells. Consistent with resistance of CIP2A^high^ cells to APR-246 *in vitro* (Fig. 1), tumor growth of CIP2A^high^ cells *in vivo* was indistinguishable between vehicle (PBS) and APR-246 treated mice (Fig. 2A). Instead, APR-246 therapy significantly decreased tumor growth of CIP2A^low^ cells (Fig. 2B). Notably, while CIP2A^low^ cells were confirmed to have almost neligible CIP2A protein expression upon transplantation (Fig. 2C), the xenograft tumors from control, or CIP2A^low^ cells were indistinguishable for their CIP2A IHC positivity at the end of the *in vivo* therapy experiment (Fig. 2D). These results indicate that CIP2A positivity in the rare population of CIP2A^low^ cells provided a strong selection advantage against APR-246 therapy.

**Figure 2.**
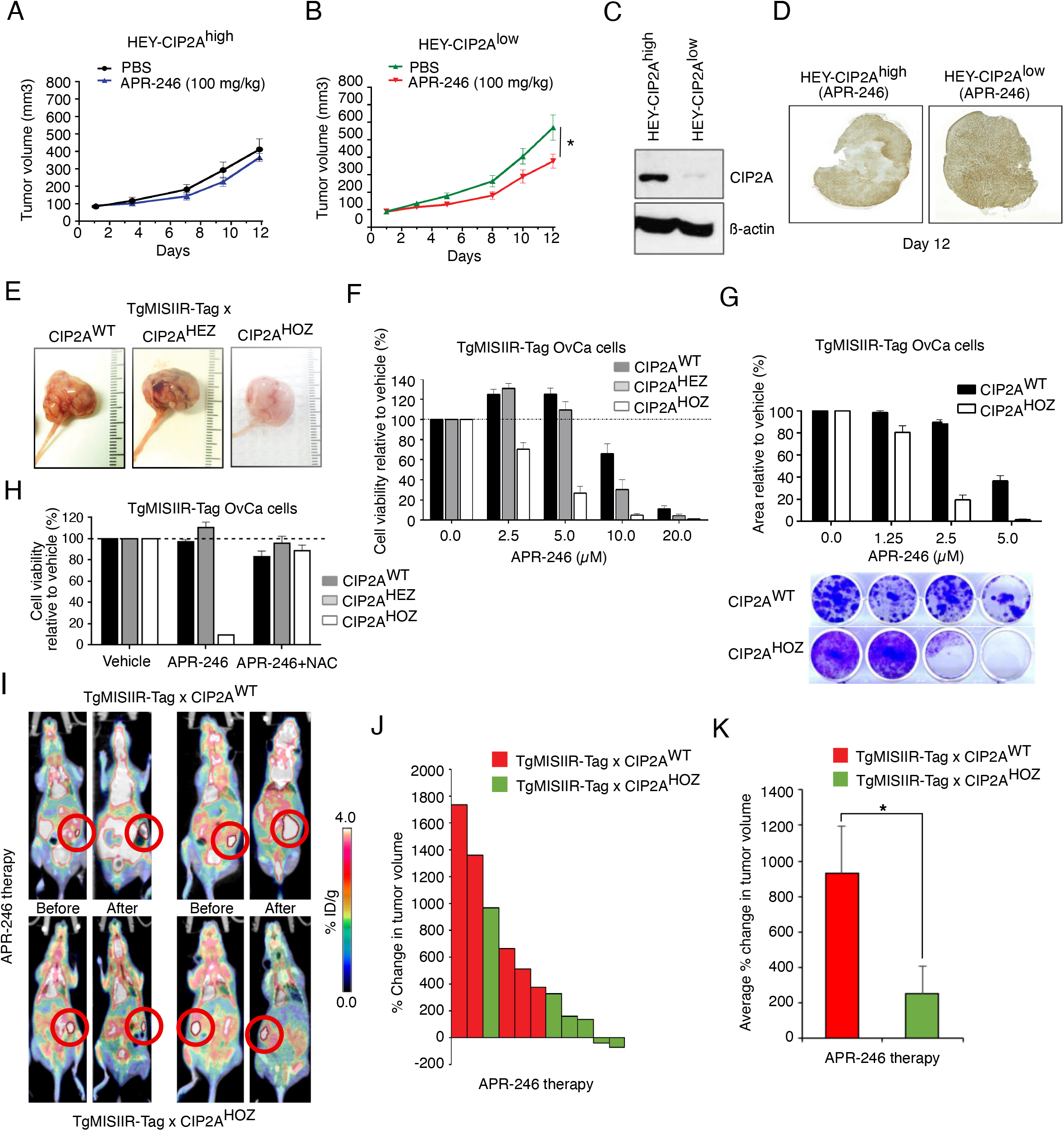
Low CIP2A expression confers OvCa cell APR-246 hypersensitivity *in vivo*. **(A,B)** Anti-tumor efficacy of APR-246 in HEY-CIP2A^high^ or HEY-CIP2A^low^ cell xenografts. Cells were injected subcutaneously in the immunocompromised mice and APR-246 treatment (5 days per week) was started after the average tumor size reached 100mm3. Shown is average tumor size from 5 mice in the group +/− S.D. * p < 0.05, t-test. **(C)** Western blot analysis of CIP2A expression levels from HEY-CIP2A^high^ or HEY-CIP2A^low^ cells before inoculation as xenografts. **(D)** CIP2A Immunohistochemistry analyses of representative end-point APR-246 treated xenograft tumors from A and B. **(E)** Representative ovarian tumors from mice with indicated genotypes. **(F&G)** APR-246 *ex vivo s*ensitivity of primary TgMISIIR-Tag murOVCAR cell lines with indicated CIP2A genotypes (combined data; n= 6 cell lines (WT & HEZ) & 4 cell lines (HOZ)). **(H)** Pre-treatment of TgMISIIR-Tag murOVCAR cells with ROS scavenger NAC rescues CIP2A^low^(HOZ) murOVCAR cells from APR-246 induced cell death. 10 μM APR-246. (**I)** PET/CT images of mice bearing TgMISIIR-Tag X CIP2A WT (upper panel) and TgMISIIR-Tag X HOZ (lower panel) tumors before and after treatment with APR-246 (100 mg/kg for 2 weeks (5 days per week)). 20-min long scans were performed 120 min post-injection of 5 MBq [18F]FDG (i.v). Tumors are highlighted with red circles. **(J)** Percentual change in metabolic active tumor volumes (MATV) between mice scanned before APR-246 treatment and two days after the last drug injection. **(K)** Average percentual change in tumor volume in response to APR-246 therapy in mice with indicated genotypes. * p < 0.05, t-test.

To further assess the *in vivo* relevance of CIP2A for APR-246 therapy response, the heterozygous and homozygous CIP2A-deficient mice (CIP2A^HEZ^ and CIP2A^HOZ^, respectively) (6) were crossed to MISIIR-TAg ovarian cancer mouse model (23). Consistent with human data that OvCa tumors may develop in CIP2A-independent manner (Table S1), we reported recently that there is no difference in OvCa tumorigenesis between MISIIR-TAg X CIP2A^WT^ and MISIIR-TAg X CIP2A^HOZ^ mice (24)(Fig. 2E). To address whether the CIP2A-deficient tumor cells from MISIIR-TAg mouse crosses yet exhibit APR-246 hypersensitivity, the OvCa cells from all three genotypes were isolated and cultured to retain their malignant characteristics as described previously (23). Fully consistent with human cell results, cells from MISIIR-TAg X CIP2A^HOZ^ mice showed dramatic hypersensitivity to APR-246 both in cell viability and colony growth assays (Fig. 2F,G). Also, similar to human cells, APR-246-elicited cell killing of MISIIR-TAg X CIP2A^HOZ^ cells was fully rescued by NAC pre-treatment (Fig. 2H).

Encouraged by these findings, we compared the *in vivo* APR-246 response of MISIIR-TAg OvCa tumors in both CIP2A genotypes by metabolic active tumor volume (MATV) measurement using PET/CT-imaging (Fig. 2I). After quantification, all but one MISIIR-TAg X CIP2A^HOZ^ tumors showed hypersensitivity to APR-246 therapy, as compared to tumors from MISIIR-TAg X CIP2A^WT^ mice (Fig. 2J). The average percentual change in tumor volume was significantly different between the genotypes (Fig. 2K). Finally, we did not observe any apparent genotype-specific differences in the weight of the mice or organs from the APR-246 treated mice, indicating that CIP2A deficiency does not result in critically limiting APR-246 hypersensitivity in the normal cells (Fig. S2A-D).

These results show that CIP2A^low^ OvCa tumors are hypersensitive to APR-246 therapy *in vivo.* However, as all the existing data related to CIP2A status in human OvCa is from diagnostic samples (8–10), it is unclear whether CIP2A^low^ tumors exists among the relapsed cases. Thereby, we surveyed CIP2A protein expression from a limited number (n=10) of available samples from HGSC OvCa ascites at disease relapse. Quantification of CIP2A protein levels demonstrated that there were clear differences between samples in CIP2A protein expression (Fig. S2E). Importantly, 4/10 of the relapsed HGSC samples (#5, #6, #8, and #10) could be clearly defined as CIP2A^low^ as compared to the rest of the tumors (Fig. S2F). These results indicate that diagnostic identification of CIP2A^low^ status in human recurrent OvCa tumors could have predictive potential for these patients regarding clinical responsiveness to APR-246 currently in clinical development (18, 19).

### Transcriptional profiling of APR-246 hypersensitive CIP2A^low^ OvCa cells

To understand the mechanistic basis of APR-246 hypersensitivity in CIP2A^low^ OvCa cells, we conducted RNA-sequencing analysis between CIP2A^high^ and CIP2A^low^ HEY cells. The parental HEY cells were included in CIP2A^high^ cohort to increase the statistical power of the gene signature enrichment analysis (GSEA), and to minimize risk that some transcriptional changes would be solely due to viral shRNA transduction. We identified 147 genes that were underexpressed, and 249 genes that were overexpressed, in CIP2A^high^ as compared to CIP2A^low^ cells (Fig. 3A, Log2 FC >1, p<0.05; Table S2). In the GSEA analysis, three transcriptional programs; Epithelial Mesenchymal Transition (EMT), TNFA signaling via NF-kB (NF-kB), and MYC targets, were significantly associated with differential gene expression profiles between CIP2A^high^ and CIP2A^low^ cells (Fig. 3B). The top ranking differentially expressed genes in these transcriptional programs are displayed in Figure 3C. Importantly, all these gene expression programs are intimately linked to OvCa pathogenesis (25–27), and MYC regulation is a hallmark for CIP2A activity in cancer cells (5). On the other hand, the identified role for CIP2A in supporting NF-kB activity in OvCa cells is consistent with recent results from breast cancer cells (28).

**Figure 3.**
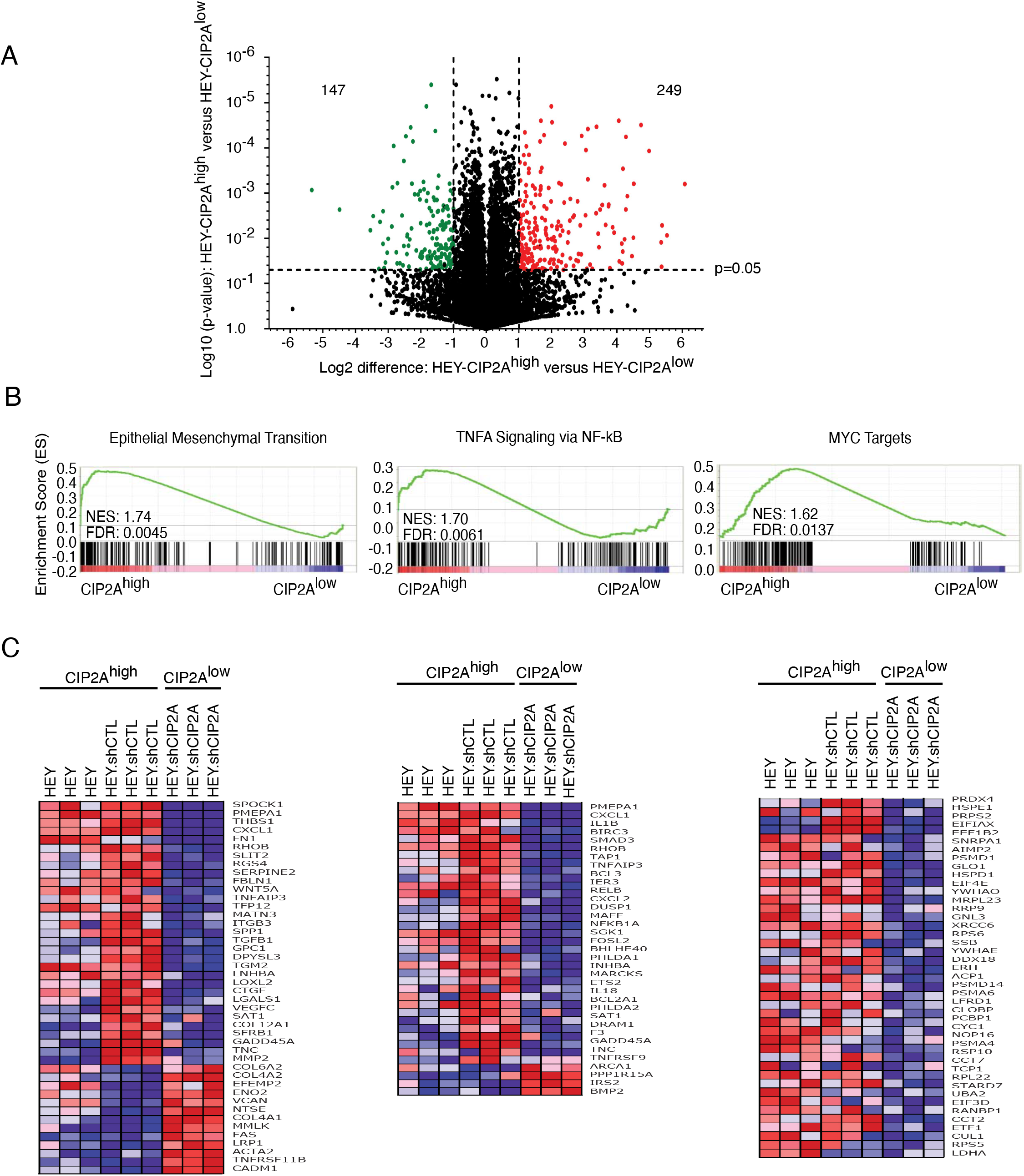
CIP2A-dependent gene expression profiles in HEY cells. **(A)** Volcano blot analysis of differentially expressed genes between HEY-CIP2A^high^ and HEY-CIP2A^low^ cells. Each dot represents one gene. Green and red dots represent significantly (Log2 <−1 or <1; p < 0.05) repressed and increased genes, respectively, in HEY-CIP2A^high^ versus HEY-CIP2A^low^ cells. **(B)** Gene Set Enrichment Analysis (GSEA) analysis of differentially expressed genes between CIP2A^high^ (includes both parental and control shRNA cells) and CIP2A^low^ HEY cells. **(C)** Heatmap presentation of the top ranking differentially expressed genes from the GSEA profiles shown in (B).

### CIP2A targets NF-kB to confer APR-246 resistance

Albeit changes in EMT, and MYC, can both contribute to drug resistance in CIP2A^high^ cells, we focused our functional validation experiments to NF-kB signaling. This was due to direct links of NF-kB to apoptosis resistance in OvCa (27), and previous data that inhibition of NF-kB inhibits cellular glutathione levels thereby potentially sensitizing cells to ROS-inducing drugs such as APR-246 (29). To begin with, we validated CIP2A-elicited regulation of selected NF-kB target genes by Q-PCR (Fig. S3A,B). Further, CIP2A^low^ HEY cells displayed significantly lower NF-kB-driven gene promoter activity (Fig. 4A). To directly assess CIP2A-mediated regulation of NF-kB, we analyzed nuclear translocation of phosphoregulated component of NF-kB complex, p65, between CIP2A^high^ and CIP2A^low^ HEY cells. CIP2A^high^ cells had significantly higher proportion of nuclear p65 than CIP2A^low^ cells in both control and TNF-α treated cells (Fig. 4B and S3C). These changes correlated with lower p65 phosphorylation in TNF-treated CIP2A^low^ cells (Fig. 4C,D). To dissect at which level of the NF-kB pathway CIP2A confers its effects, we studied NF-kB promoter activity in combination with overexpression of the p65 upstream kinase MEKK3 (30). MEKK3 overexpression strongly induced NF-kB promoter activity in CIP2A^high^ cells, but this was blunted in CIP2A^low^ cells (Fig. 4E; lane 3 vs. 4). However, CIP2A inhibition was able to blunt NF-kB activity also in cells overexpressing MEK mutant with non-dephoshorylatable serine 250 and threonine 516 (MEKK3^S250D/T516D^) (Fig. 4E; lane 7 vs. 8). These findings together with CIP2A effects on p65 phosphorylation (Fig. 4C,D), support the conclusions that CIP2A promotes NF-kB activity downstream of activated MEKK3.

**Figure 4.**
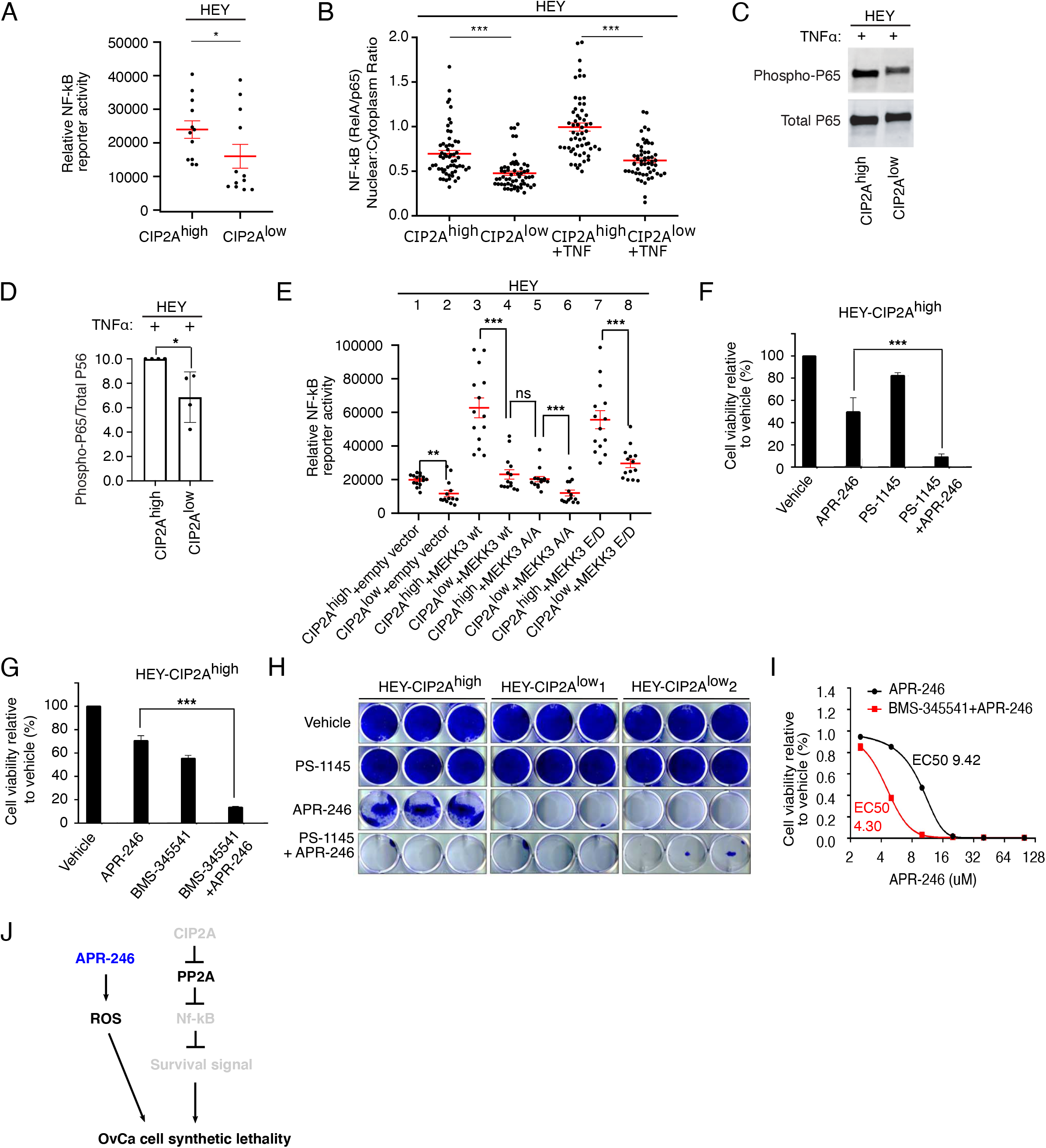
CIP2A promotes NF-kB activity in APR-246 insensitive OvCa cells. **(A)** Relative NF-kB luciferase reporter activity in HEY-CIP2A^high^ and HEY-CIP2A^low^ cells. Shown is mean + S.E.M. * p<0.05, Mann-Whitney test. **(B)** Quantification of p65 signal intensity ratio (Nuclear/Cytoplasm) in HEY-CIP2A^high^ or HEY-CIP2A^low^ cells with or without TNF-alpha treatement. Shown is mean + S.E.M. *** p<0.001, Mann whitney test. **(C)** Western blot analysis of phospho-P65 and total p65 from TNF-alpha treated HEY-CIP2A^high^ or HEY-CIP2A^low^ cells. **(D)** Quantification of relative p65 phosphorylation from (C). n=4. *** p<0.05, Mann-Whitney test. **(E)** Relative NF-kB luciferase reporter activity in HEY-CIP2A^high^ or HEY-CIP2A^low^ cells with either empty vector, MEKK3 WT, MEKK3 T516A/S250A, or MEKK3 T5163/S250D overexpression. Shown is mean + S.E.M. *** p<0.001, Mann whitney test. **(F,G)** Relative cell viability of HEY-CIP2A^high^ treated with APR-246 alone, or with IKK inhibitors PS-1145 or BMS-345541 alone, and their combinations. *** p<0.001, t-test. **(H)** Colony Growth assay of HEY-CIP2A^high^, HEY-CIP2A^low1^ and HEY-CIP2A^low2^ treated with either vehicle, PS-1145 alone, APR-246 alone or PS-1145 + ARP-246. **I)** Relative cell viability in patient-derived HGSC cell line OC002 treated with either APR-246 alone or BMS-345541 + APR-246. EC50 values for APR-246 in each condition are indicated next to concentration curve. **J)** Schematic model of mechanistic basis of CIP2A-mediated APR-246 resistance in OvCa cells. Grey colour denotes for situation where the target is inhibited.

To address whether CIP2A-driven NF-kB activity functionally confers APR-246 resistance, we tested whether similar synergy that was observed between CIP2A inhibition and APR-246, could be recapitulated by co-treatment of CIP2A^high^ cells with APR-246 and small molecule inhibitors of NF-kB. As a result, all three tested NF-kB inhibitors, each with different mode of action, potentiated the effects of APR-246 in inhibition of cell viability in CIP2A^high^ HEY cells (Fig. 4F,G, S3D). These results were substantiated by colony growth assays including two independent CIP2A^low^ HEY cell clones with different CIP2A shRNAs. With the chosen dose, NF-kB inhibitor PS-1145 did not have any notable effect on either CIP2A^high^ or CIP2A^low^ cells, but in combination with APR-246 it induced similar synthetic lethal phenotype that was observed with APR-246 in CIP2A^low^ cells (Fig. 4H). Finally, the combined action of APR-246 and NF-kB inhibition was validated in patient-derived OvCa cell line OC002 derived from a patient with disseminated disease (Fig. 4I)(11). These results indicate that inhibition of NF-kB activity mediates APR-246 sensitivity in CIP2A^low^ OvCa cells (Fig. 4J).

During relapse from chemotherapy, the OvCa cells have exhausted their capacity to respond to DNA-damaging agents (1), but might yet be vulnerable to other therapies in a subtype specific manner (2). Our results collectively identify CIP2A as a context-dependent oncoprotein in OvCa. It is dispensable for both human and mouse OvCa tumorigenesis, but associates with more aggressive disease (8)(Table S1), and drives resistance to APR-246 therapy. Together with our analysis from a limited number of available human relapse samples, these data indicate that OvCa tumors with low CIP2A expression constitute a minor, but yet clinically relevant novel human OvCa subtype. Combined with recently demonstrated role for CIP2A in confining therapy response for dozens of commonly used cancer drugs in other cancer cell types (12, 13), our results encourage further screening of CIP2A^low^ OvCa cell models against larger drug libraries to identify, in addition to APR-246, other drugs to be tested for the treatment of CIP2A^low^ OvCa subtype patients.

Current data indicate that APR-246 kills cancer cells via multiple mechanisms (16, 17, 20–22). Our data about ROS-dependent, but most likely TP53-independent mechanism of OvCa cell killing by APR-246 is directly supported by recently published work (17, 21). This is potentially clinically important finding as it indicates that *TP53* status would not dictate the cell killing activity of APR-246 in CIP2A^low^ OvCa subtype tumors. APR-246 has been tested in two OvCa clinical trials (NCT02098343, NCT03268382) but no results are publicly available. Currently, APR-246 is studied in clinical trials in AML and myelodysplastic syndromes (18), and in various other solid cancer types (https://www.clinicaltrials.gov). Similar to OvCa, also among these cancer types there is a significant number of patients with CIP2A^low^ subtype (4, 7). Therefore, and acknowledging the role of NF-kB activity in regulation of cellular buffering capacity against ROS (29), it would be very interesting to examine CIP2A expression and NF-kB pathway activity, from the clinical trial patient samples from these past and ongoing APR-246 trials. By these means the presented results could support future ARP-246 clinical trials in better predicting the potential responders, and thus establish a future patient stratification strategy for clinical use of APR-246. In addition, our results position CIP2A as an APR-246 combination therapy target for ovarian cancer.

## Acknowledgements

We are very grateful to APREA Therapeutics for APR-246 compound. Professor Caj Haglund, and Dr. Ralf Butzow are acknowledged for their contributions to the original CIP2A expression analysis from the OvCa cohort. This study used Turku Bioscience Centre core services by Finnish Functional Genomics Centre, and Cell Imaging and Cytometry, funded by University of Turku and Åbo Akademi University and Biocenter Finland. The work was funded by Sigrid Juselius Foundation (JW), Finnish Cancer Foundation (JW), Helsinki University Central Hospital Research Funds (AR) and Finska Läkaresällskapet (AR). Fox Chase Cancer Center (FCCC) Core Grant NCI P30 CA006927 (DCC) and generous donations from the Dubrow Fund and the Bucks County Board of Associates and the Mainline Board of Associates (DCC), and acknowledges the FCCC Laboratory Animal Facility for the husbandry and exportation of the TgMISIIR-TAg mice used in this study. TDL was funded as a FICAN Cancer Researcher for the Finnish Cancer Institute and by the Finnish Cultural Foundation.

